# Interpersonal Synchrony and Implicit Identification

**DOI:** 10.1101/2025.04.16.649093

**Authors:** Manisha Biswas, Marcel Brass

## Abstract

The transformative effects of interpersonal synchrony on reshaping the relationship between the self and the group by fostering self-other blurring are well-documented. However, these findings largely rely on self-reported measures, leaving aspects of the self inaccessible to explicit measures unexplored. We combined the Implicit Association Task (IAT) with conventional explicit evaluations of self-other blurring to examine how experiences of synchronous movement may also swiftly shape implicit measures indicative of self-other blurring. In three online experiments participants experienced synchronous movement as opposed to no movement (Experiment 1) or unpredictable movement (Experiment 2) or asynchronous movement (Experiment 3) with virtually presented agents. Our data indicate that synchronous movement led to higher levels of self-other blurring on the Inclusion of Other in Self (IOS) scale and implicit self-concept evaluations on the IAT. Participants demonstrated a preference for associating self-related words with images of the synchronous group, while associating other-related words with images from the no-movement, unpredictable, and asynchronous movement groups. This suggests that experiencing synchronous movement with a group influences implicit semantic self-associations. These findings suggest that interpersonal synchrony may alter self-concept at both an embodied and semantic level, extending beyond conscious affiliation. Our combined methodology offers a deeper insight into the impact of synchronous experiences on self-constructs and social relationships.

## Introduction

### Self-constructs and Collective Identity

An individual’s self-construct is not solely defined by personal characteristics, it is also moulded by an awareness of the group memberships (Mattingly et al., 2020; Tajfel et al., 1971, 1986). Self-constructs consist of an individual’s interpersonal knowledge, shaping how they engage with others and the broader social world (McConnell, 2011). They are flexible and dynamic, adapting to the nuances of current social situations (Markus & Wurf, 1987). It has been previously demonstrated that group membership can alter the size, structure and diversity of an individual’s self-concept as the boundaries between the self and the group become blurred (Mattingly et al., 2020). Such shifts in self-constructs influence cognitive representations of the self and group alignment (Griffin & Bartholomew, 1994; Mikulincer et al., 2003; Smith et al., 1999).

Ritualised experiences of collective synchronous movement facilitate one’s induction into a group and serve to maintain cohesive groups (Cirelli, 2018; Cirelli et al., 2016; Whitehouse & Lanman, 2014). A growing body of research suggests that the experience of interpersonal synchrony serves as a key marker of group membership and is associated with increased social closeness and pro-social attitudes (Hove & Risen, 2009; Mogan et al., 2017; Reddish et al., 2013; Valdesolo & DeSteno, 2011). Self-other blurring serving as a key mechanistic account for the prosocial effects generated through synchrony (Aron et al., 2013; Hove & Risen, 2009; Launay et al., 2016; Mazzurega et al., 2011; Paladino et al., 2010; Swann et al., 2012). This suggests that interpersonal synchrony not only creates greater group affiliation but also affects the individual at a more fundamental level via change in self-constructs (Reddish et al., 2020)

When individuals engage in synchronous actions, the boundaries between their own identity and that of synchronous bodies become blurred (Decety & Sommerville, 2003; Hove & Risen, 2009; Smith, 2008; Swann et al., 2012). Drawing on the rubber hand illusion, researchers argue that witnessing someone else perform the same actions as us leads to a sense of shared identity and increased sense of control (Launay et al., 2016). In 2020, Reddish et. al. argued that the self is a complex construct, and interpersonal synchrony influences one of its fundamental dimensions – sense of agency. Sense of agency refers to the feeling that we have caused some action or event in the world. It depends on temporal contiguity between action and effect (Pacherie, 2014; Synofzik et al., 2008; Wegner et al., 2004). When self-action is paired closely in time with other-action, like in the case of rhythmic interpersonal synchrony, participants are likely to develop an increased sense of agency because they feel as if they control others (Pacherie, 2012; Reddish et al., 2020). This is also known as extended self-agency or the feeling that one has agency over the actions of another person’s actions. However, the relationship between interpersonal synchrony is not straight-forward as while it might increase self-agency it is also associated with an increase in other-agency. Reddish et. al. suggest that this might be due to the collaborative nature of interpersonal synchrony.

### Limitations of Explicit Measures in Assessing Self-Other Blurring

The predominant method employed so far in support of self-other blurring account of synchrony is a single item pictorial self-report scale known as the Inclusion of the Other in Self scale, or IOS (Aron et al., 1992; for meta-analysis, Rennung & Göritz, 2016). The IOS scale directly prompts participants to assess the extent of overlap between themselves and a group through images of gradually overlapping circles (see Fig. 1). In practice, using the IOS scale involves participants rating the degree of overlap immediately after experiencing synchronous (as compared to asynchronous) movements with an agent(s). As such, the measure has been criticised for being susceptible to demand characteristics because it is relatively easy for participants to discern the responses researchers are seeking (Orne, 1959, 2017), including by its developers (Aron et al., 1991; Aron & Fraley, 1999). Numerous iterations of this scale have evolved over time (Jiménez et al., 2016), with documented modifications such as the added option of complete overlap (Schubert & Otten, 2002; Swann et al., 2009) and the introduction of a verbal accompaniment (Gómez et al., 2011), but pre-existing modifications still exclusively target the explicit dimension of self-other blurring.

**Figure 1.**
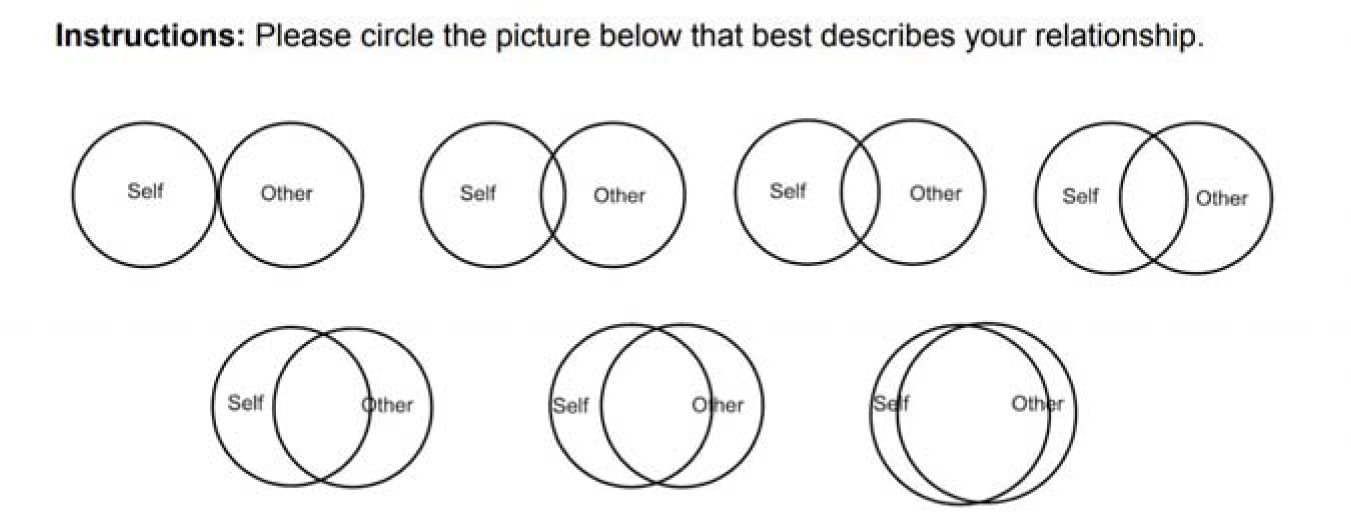
Reproduction of the IOS (Inclusion of Other in Self) Scale from Aron et. al (1992).

Responses on the IOS can be interpreted as attitudes rather than traits as the degree of self-other overlap with one group is not correlated with the tendency to report higher degrees of self-other overlap with another group (Swann et al., 2012). According to the Dual Attitude Model, attitudes toward a group comprise of both explicit and implicit dimensions (Wilson et al., 2000). Self-report measures like the IOS scale are conceptually limited because they are primarily designed to capture explicit attitudes. Explicit attitudes can be consciously reported and controlled and are often measured using self-report, whereas implicit attitudes cannot be consciously accessed or controlled. They are inferred using indirect measures such as the me/not me categorisation task (Aron et al., 1991) or the IAT (Greenwald & Lai, 2019). Researchers have found that implicit attitudes are pervasive in many forms of social cognition (for review: Greenwald & Lai, 2019) and there is mounting evidence that self-concepts operate implicitly (Asendorpf et al., 2002; Briñol et al., 2006; Greenwald & Banaji, 1995; Hetts et al., 1999; Mattingly et al., 2020). By examining implicit semantic associations with self-constructs, tools like the IAT elucidate a distinct dimension of self-other blurring (Bargh et al., 1992; Greenwald & Banaji, 1995), and in contexts involving shared group identity, implicit measures are often necessary to detect intergroup biases (Greenwald et al., 1998; Greenwald & Banaji, 1995). The potential for interpersonal synchrony to alter implicit self-associations is largely unexplored.

It is crucial to acknowledge the ongoing debate regarding the explicit-implicit terminology (Fazio & Olson, 2003; Gawronski, 2019; Greenwald & Banaji, 1995; Schimmack, 2021). However, we believe that when applied appropriately, the explicit-implicit construct can serve as a valuable foundation for further investigation, particularly when exploring explicit attitudes alongside implicit attitudes as they both offer insights into separate facets of the underlying psychological constructs (Gawronski, 2018). To address this debate, we ground our differentiation in Gawronski’s (2018) theoretical framework by emphasising the observable characteristics of the measured response. We employ the terms *explicit and implicit evaluations* to specify that we are referring to behavioural evaluative responses. Our theoretical emphasis on the distinction stems from the finding that explicit and implicit evaluations independently predict behaviours (Dovidio et al., 2002; Kurdi et al., 2019) and choices (Galdi et al., 2008). Researchers sometimes find double-dissociation patterns when studying implicit and explicit evaluations together, such that, implicit evaluations are better at predicting spontaneous behaviour whereas explicit evaluations are better at predicting deliberate behaviour (Friese et al., 2008; Perugini et al., 2010). For example, implicit evaluations are more consistently predictive of nonverbal behaviour during interracial interactions, whereas explicit evaluations show a stronger predictive association with the content of verbal responses in these interactions (Dovidio et al., 2002). Although the motivation for socially desirable responding may not be as strong as in interracial prejudice paradigms, it is theoretically important for the field of interpersonal synchrony to determine whether implicit and explicit self-other blurring dissociate or converge.

### Advancing Research with Implicit Measures of Self-Other Blurring

Given the differences between explicit and implicit evaluations, investigating the implicit effects of synchrony-induced self-other blurring is crucial. We argue that interpersonal synchrony affects the self at different levels of psychological processing. The IOS Scale can only account for the embodied explicit level, but some forms of group affiliation can only be inferred at an implicit semantic level. Additionally, implicit evaluations were traditionally described as slow to adapt and resistant to transient interventions (Petty et al., 1997). Some researchers argued that differences in implicit evaluations could only be elicited by invoking membership to long-standing groups (Rydell & McConnell, 2006). However, several recent studies have found that minimally assigned groups in a lab setting can also elicit implicit intergroup bias favouring in-group members (Otten & Moskowitz, 2000; Xiao & Van Bavel, 2019), indicating that implicit evaluations demonstrate a notable degree of flexibility and can be influenced by varying group (Gawronski, 2018). Moreover, mental representation of the self are known to be flexible and context-sensitive constructions that change in accordance with social situations and current goals (Smith, 1998; Smith et al., 1999). It remains to be empirically assessed whether interpersonal synchrony induced minimal groups can shift implicit attitudes.

Prior research shows that as we form closer relationships with others, our self-concept becomes blurred not only in the explicit sense captured by the IOS Scale but also in an implicit semantic dimension, where distinguishing between self and other becomes increasingly difficult (Slotter & Gardner, 2009, 2012b, 2012a; Smith et al., 1999). Majority of experimental evidence supporting the idea of interpersonal relationship-induced changes in implicit self-construct primarily stems from research on romantic relationships. Engaging in romantic relationships often prompts individuals to integrate their partners into their self-concept, leading to a shift from a self-centred mindset to the adoption of a collective mindset (Agnew et al., 1998; Andersen & Chen, 2002; Rusbult & Agnew, 2010). These experiments focus primarily on a known *‘other’*, for example, a romantic partner.

We propose that a comparable process may occur in experiments that induce interpersonal synchrony with a *‘exchangeable’* other or interaction partner, as the mechanisms identified for maintaining romantic relationships, such as prosocial tendencies and a greater willingness to make sacrifices (Agnew & Le, 2015; Smith et al., 1999; Van Lange et al., 1997), have also been recognised by researchers studying interpersonal synchrony (Whitehouse & Lanman, 2014). When we engage in synchronous movement with other agents, it initiates a cognitive restructuring and/or re-evaluation of one’s self-concept by acquiring new or enhancing existing content, potentially affecting both explicit and implicit self-constructs (Gordon & Luo, 2011).

Using the IAT to assess self-other blurring serves a twofold purpose (1) it gives us access to the semantic level of group affiliation and, (2) it increases the credibility of the self-other blurring account of synchrony. Latency-based categorization tasks, such as the IAT assess the strength of associations between the self and various concepts stored in memory, including group-related associations (Kurdi et al., 2019; Perkins & Forehand, 2006). While the IOS scale captures a consciously accessible and embodied sense of closeness between the self and others, it may not fully reflect automatic, semantic changes in the self-concept. Interpersonal synchrony may exert its effects not only at the level of felt affiliation, but also through changes in non-conscious identity structure. For this reason, we argue that it is necessary to distinguish between embodied self–other overlap, which is reported directly via the IOS, and semantic self–other association, which may be better captured through implicit methods such as the IAT (Perkins & Forehand, 2006; Swanson et al., 2001). The IAT allows us to assess the extent to which the concept of the self becomes cognitively linked with a synchronised group at a semantic level. These associations operate below the level of conscious awareness and are not easily accessible through introspection. Thus, employing the IAT enables us to examine whether interpersonal synchrony reshapes implicit self–group associations—offering a more nuanced understanding of how identity may be transformed through shared action.

Moreover, although the IOS is an extensively adopted and well-validated measure, it is potentially vulnerable to demand characteristics (Aron et al., 1991; Aron & Fraley, 1999). The IAT mitigates this issue by relying on speeded responses and less transparent stimulus–response pairings, thereby reducing the likelihood of hypothesis-driven responding as response latencies are largely immune to self-presentational biases (Kurdi et al., 2019; Paulhus, 1984; Slotter & Gardner, 2009). Including both measures allows us to triangulate effects and identify the extent to which observed self–other overlap is robust across different levels of awareness and susceptibility to demand.

## Current Study

### Avatar-mediated virtual synchrony

While most previous studies on synchrony have involved in-person, physical movement, the present work uses avatar-mediated synchrony to investigate how temporal alignment with others may affect self-concept. Prior work has shown that even in virtual settings, synchrony can produce similar effects on explicit self-other blurring (Biswas & Brass, 2024). While avatar-mediated synchrony has not yet been directly compared to real-life synchrony in the literature, existing evidence suggests it produces similar effects on the Inclusion of Other in the Self (IOS) scale—the most widely used and replicated measure of self–other overlap in interpersonal synchrony research. This convergence on a canonical outcome provides validity to the claim that avatar-mediated online interpersonal synchrony evokes similar processes to in-person synchrony paradigms. Although participants do not perform full-body movements themselves, they control an avatar whose actions are temporally synchronised with those of other agents. This paradigm preserves core components of synchrony—shared timing, perceived extended agency, and coordinated action—while allowing for precise manipulation of experimental conditions.

A notable benefit of online paradigms is the ability to control and systematically adjust group size in synchrony research. Much of the anthropological work that inspired research on interpersonal synchrony emphasised its role in large-scale collective rituals, where acting with a group produces qualitatively different experiences than coordinating with a single partner (Collins, 2004; Durkheim & Fields, 1995; Rimé & Páez, 2023). Despite this, empirical research has been limited to dyadic interactions (Mogan et al., 2017), due to the logistical difficulties of standardising collective synchrony without introducing session-level variability. This focus on dyads overlooks the fact that group size can moderate the psychological outcomes of synchrony (Launay et al., 2016; Tarr et al., 2014, 2015).Virtual environments now offer a powerful tool to overcome these constraints, allowing researchers to study synchrony in group contexts with greater control and flexibility.

We conducted three experiments to explore the effects of avatar-mediated synchrony on both explicit and implicit self-other blurring. This setup involved creating an experience of synchrony that could be modulated in real-time among a group of virtually presented avatars (see Fig. 2). In our experimental design, participants controlled an avatar that moved in response to their key presses. They were positioned opposite two groups of agents whose movements varied depending on the experimental condition, In Experiment 1, we compared a synchronous movement condition to a no-movement control condition to determine whether synchrony influenced both explicit and implicit self-evaluations. In Experiment 2, we introduced an unpredictable movement control condition, where the agents moved independently and without coordination. Finally, in Experiment 3, we compared synchronous movement to a traditional asynchrony condition. Agents in the control condition made the same movement but at an altered phase (see panel 3, Fig 2). These experiments allowed us to explore the effects of synchrony across varied control conditions.

**Figure 2.**
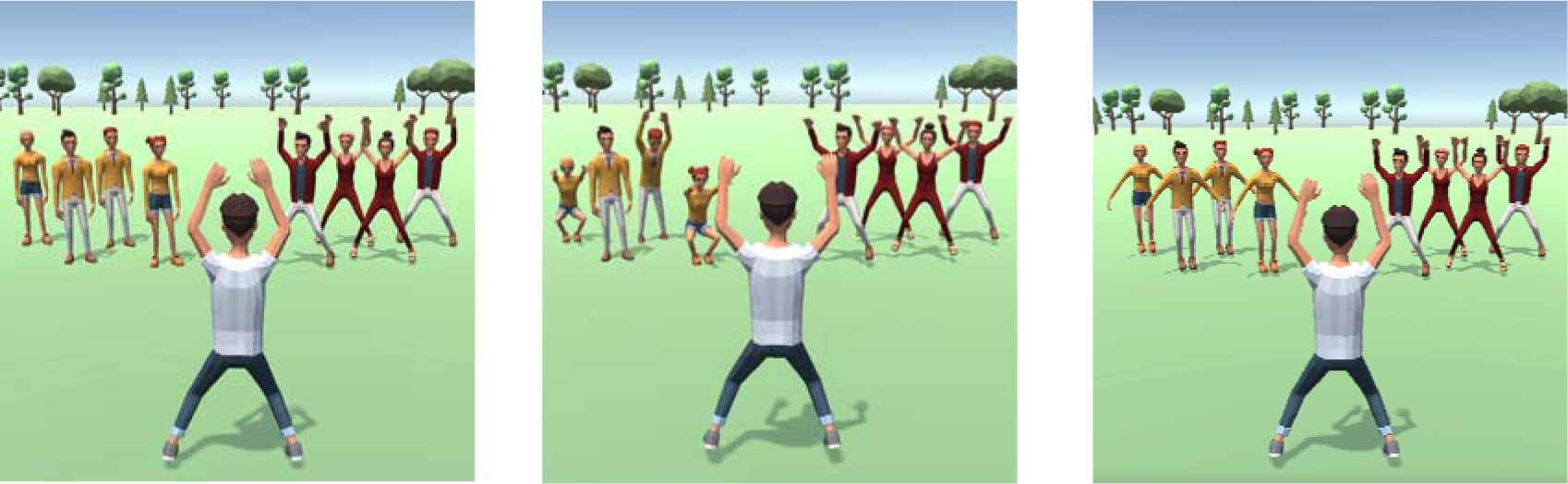
Synchrony Manipulation in Experiment 1: Synchronous movement versus No movement (left cell), Experiment 2: Synchronous movement versus Unpredictable movement (middle cell) and Experiment 3: Synchronous movement versus Asynchronous movement

### Self-Other IAT

To test whether these experiences of interpersonal movement synchrony with a group can influence implicit evaluations of the self, we use an adapted version of the Self-Other Implicit Association Task or the IAT. The self-other IAT is a derivative of the classical IAT task (Greenwald et al., 1998), a procedure that assesses the strength of an association between a target concept and an attribute dimension, by considering the latency with which participants can employ two response keys when each has been assigned a dual meaning. The Self-Other IAT is an IAT variant, often used to study self-esteem by using self-other as target concepts and positive-negative as the attribute dimensions (Greenwald & Banaji, 1995). We adapted this procedure to our experimental set-up by replacing the target concept with pictures of the groups presented in our manipulation and replacing the attribute dimension with self and other-words to discern which group was implicitly constructed as closer to the self. Our hypothesis was that the experience of synchronous movement with one of the groups would shift self-concepts such that the synchronous group would be incorporated into the self.

### Agency Ratings

To investigate sense of control over self–avatar, synchronous group and control group. We asked participants in Experiments 2 and 3 about their sense of control over their avatar, the synchronous group, and the control group. This allowed us to assess differences in perceived control across synchronous, unpredictable, and asynchronous movement conditions, examining how movement presence, predictability, and temporal lag shaped agency.

### Experiment 1

#### Methods

Methods and the data analysis plan were pre-registered (https://aspredicted.org/RLP_TVW). We determined our sample size in advance of any data collection, and we report all data exclusions, all manipulations, and all measures in the study. The experiment was programmed on Unity (ver. 2022.3) using the UXF 2.0 package (Brookes et al., 2020) and administered online using JATOS (ver. 3.8.1).

#### Participants

We collected the data of 60 participants (43% female, 1.8% non-binary, M_age_ = 36, SD_age_ = 10) from Prolific. 7 participants were excluded based on pre-registered criteria. In particular because they failed the attention check (see supplementary materials). All participants were naïve to the aims of the experiment and compensated at a rate of 6 pounds/hour for their time. Participants provided informed consent before they began the experiment. The experimental procedures were approved by the ethical review board at the Institute of Psychology, Humboldt University of Berlin and were conducted in accordance with the American Psychological Association’s Ethical Principles in the Conduct of Research with Human Participants.

### Procedure

#### Synchrony Manipulation

Computer-mediated environments, resembling video games, were previously shown to have significant influence on self-concepts (Klimmt et al., 2010). In an online desktop experiment, participants were instructed to control their avatar using the space key, and to press the key when they heard an audio cue. The audio cue was presented 15 times per block at a rhythm of 60 bpm. Each time the participant pressed the spacebar, their avatar moved in a jumping jacks motion (see Fig. 2). This animation took approximately 900 *ms* to complete.

Participants were exposed to two groups of virtual avatars facing their avatar, one on the left and the other on the right (see Fig. 2). Each group of avatars was composed of two male-presenting and two female-presenting agents, both groups faced the participant’s avatar. For half of the participants the left group moved in synchrony with the participants’ avatar and for the other half, the right group moved in synchrony while the other group showed no movement (see Fig. 2 left cell). To ensure that the participants felt the highest degree of control over their avatar, the first 5 times the audio cue was presented, only their avatar moved. After the 6th beat, the synchronous group would show the same movement as the participants’ avatar.

After undergoing the synchrony manipulation, participants engaged in the implicit association task. Our task was designed in accordance with the well-replicated utilised paradigm outlined by Greenwald et al., (1998). It comprised the sections displayed in table 1. During the first categorisation phase, participants were shown images of either the synchronous group or the no-movement group. Adjacent to these images, labels appeared on the left and right sides instructing participants to press the *“e”* key if the image belonged to the synchronous group and the *“i”* key if it belonged to the no movement group (or vice versa, depending on the order of the compatible or incompatible blocks, see table 1). In a subsequent section, participants encountered self-related words (e.g., *I*, *me*, *mine*, *my*, *self*) and other-related words (e.g., *other*, *their*, *theirs*, *them*, *they*). Their task was to rapidly categorise these words, pressing the *“e”* key for self-related words and the *“i”* key for other-related words.

**Table 1.** Design of the implicit association task compatible first.

| Block | No. of trials | Function | Right Key Assignment | Left Key Assignment |
| --- | --- | --- | --- | --- |
| 1 | 20 | Practice: Image | Sync Group Picture | Control Group Picture |
| 2 | 20 | Practice: Self-Other | Self-words | Other-words |
| 3 | 60 | Test: Combination 1 | Sync + Self-words | Control + Other-words |
| 4 | 20 | Practice: Image Reverse Mapping | Control Picture | Sync Picture |
| 5 | 60 | Test: Combination 2 | Control + Self-words | Sync + Other-words |

**Note.** For participants that faced incompatible blocks first, block 1 was replaced with block 4 and block 3 was replaced with block 5

#### Implicit Association Task

The subsequent phase presented participants with their first combined block, in which they encountered either the compatible or incompatible blocks. In the compatible blocks, they were instructed to press the *“e”* key for self-words or synchronous group image presentations and the *“i”* key for other-words and non-movement group image presentations. In the incompatible blocks, the attribute dimension was reversed: participants pressed the *“e”* key for self-words and no movement group images, and the *“i”* key for other-words and synchronous group image presentations. Following this, participants underwent a block in which the key mapping for synchronous group and no movement group image presentations was reversed. Subsequently, they encountered another block of combined trials. If they had initially faced a compatible block, they now encountered an incompatible one, and vice versa.

#### Post-task questionnaires: IOS Scale and Attention Check

Participants were then presented with images of the two groups asked to provide a score on the IOS scale for both. They were also asked to select which group moved in synchrony with them as an attention check.

## Data Processing and Analysis plan

We used our pre-registered subject and trial exclusion criteria to clean our data (see supplementary materials) and calculated the D measure, a procedure to calculate the differences between the compatible and incompatible blocks in an IAT (Greenwald et al., 2003).

## Results

### Self-Other Blurring

We compared IOS Scale scores reported for the Synchronous Movement and No Movement groups. IOS scores were not normally distributed according to the Shapiro-Wilk test (W = 0.938, p = .009). Hence, we report Wilcoxon signed-rank test values for this effect. Results showed that IOS scores were significantly higher for the Synchronous Movement group (M = 4.22, SD = 1.98) as compared to the No Movement group (M= 1.68, SD = 1.23) (W = 938, p <.001) in a directed t-test (Synchronous Movement > No Movement) indicating that the explicit self-other blurring was significantly higher for the synchronous movement group.

**Figure 3.**
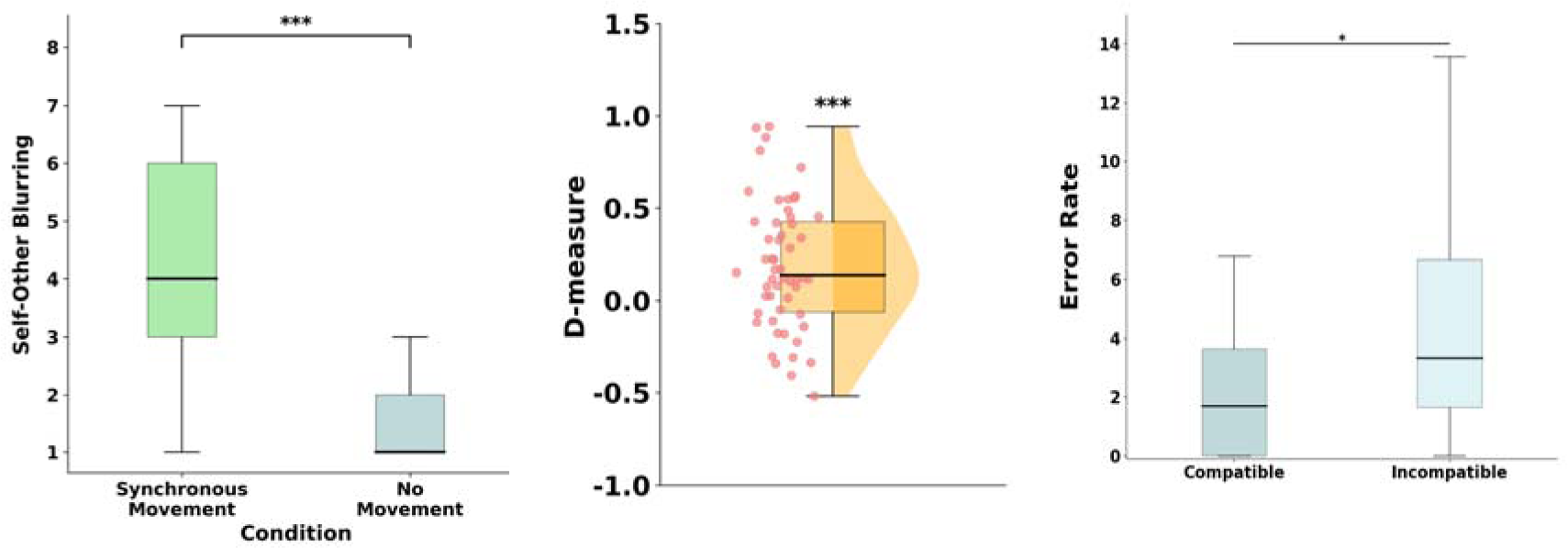
Comparison of IOS Scores for Synchrony vs. No Movement Group. Cell 1: Inclusion of Other in the Self (IOS) scores for the Synchrony and No Movement groups. Cell 2: Implicit Association Test (IAT) D measure between the Synchrony and No Movement groups. Cell 3: Error rates for the Synchrony and No Movement groups

### IAT: D measure

We predicted that participants would develop an implicit association between their self-construct and synchronous movement group that would lead them to associate self-words with images of the synchronous group and other-words with images of the no movement group. To test our hypothesis, response latencies for the blocks of the combined task (Blocks 3, 4, 6, and 7) were recorded and analysed according to the improved IAT scoring algorithm (Greenwald et al., 2003). We computed the *D* score by dividing the differences in the latency scores between the incompatible and compatible IAT blocks by the standard deviation of all the correct trials in the combined blocks. A positive *D* score represents a faster response to compatible pairings (self + synchronous movement group and other + no movement group) than to incompatible pairings (self + no movement group and other + synchronous movement group). The *D* score (*M* = 0.410, SD = 0.455) was significantly greater than zero (*t_(52)_*_=_ 6.566, *p* < .001, 95% CI [0.579, 1.219]), indicating that the participants tended to react faster to compatible pairings than to incompatible pairings. This result suggested that the synchronous movement group was associated with self-words more strongly than with other-words and that the no movement group was associated with other-words more strongly than with self-words.

### IAT: Error Rate

We compared the error rate across the compatible and incompatible blocks. Error rates were not normally distributed according to the Shapiro-Wilk test (W = 0.630 p < .001). Hence, we report Wilcoxon signed-rank test values for this effect. Results showed that error rates were significantly lower in the compatible blocks (M = 2.74, SD = 3.3) as compared to the incompatible (M= 4.8, SD = 7.2) (W = 379, p =0.031) in a directed t-test (Compatible < Incompatible).

**Figure 4.**
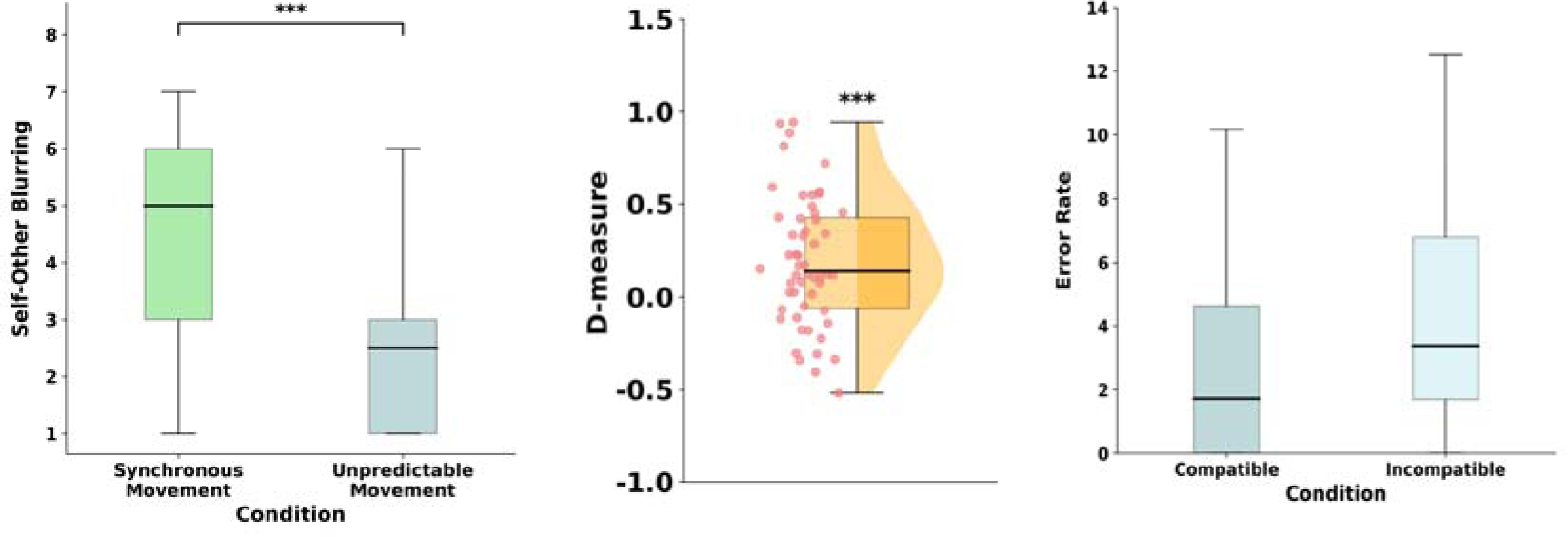
Comparison of IOS Scores for Synchrony vs. Unpredictable Movement Group. Cell 1: Inclusion of Other in the Self (IOS) scores for the Synchrony and Unpredictable Movement groups. Cell 2: Implicit Association Test (IAT) D measure between the Synchrony and Unpredictable Movement groups. Cell 3: Error rates for the Synchrony and Unpredictable Movement groups.

### Correlation between Implicit and Explicit Measures

To establish whether there was a relationship between self-other blurring and D measure, we conducted a correlation analysis between the IOS difference and D measure across participants. We were unable to find a significant correlation (r = 0.054, 95% CI [-0.219, 0.320], p = 0.701).

## Interim Discussion

The results of Experiment 1 suggest a significant impact of synchronous movement on the explicit self-other blurring, as indicated by higher IOS scores in the Synchronous Movement group compared to the No Movement group. Additionally, the IAT D measure demonstrated a strong implicit association between the self and the synchronous movement group, with participants responding faster to compatible pairings (self + synchronous movement group, other + no movement group). The lower error rates in the compatible blocks further support the reliability of this association. In a follow-up experiment we introduced a new control condition of unpredictable movement to replace no movement (see Fig. 2, second cell). This adjustment was made to address concerns raised from Experiment 1, specifically regarding the presence of movement, which may have attracted more attention and potentially influenced the observed effects. We also added a question regarding direct control of the participants’ avatar, the synchrony group and the unpredictable movement group.

### Experiment 2

#### Methods

Methods and the data analysis plan were pre-registered (https://aspredicted.org/FCX_77L)The methods used were identical to Experiment 1, but the control condition was switched from a group that showed no movement to a group that showed unpredictable movement. We determined our sample size in advance of any data collection, and we report all data exclusions, all manipulations, and all measures in the study. The experimental procedures were approved by the ethical review board at the Institute of Psychology, Humboldt University of Berlin and were conducted in accordance with the American Psychological Association’s Ethical Principles in the Conduct of Research with Human Participants.

#### Participants

We collected the data of 60 participants (40% female, M_age_ = 34.8, SD_age_ = 9.7) from Prolific. 5 participants were excluded based on our pre-registration criteria because they failed the attention check, additionally 1 more participant was removed because they did not provide a response to the avatar control question. All participants were naïve to the aims of the experiment and compensated at a rate of 6 pounds/hour for their time. Participants provided informed consent before they began the experiment. The experiment was approved by the ethical review board at the Institute of Psychology, Humboldt University of Berlin and was conducted in accordance with the directives laid out in the Helsinki Declaration of Human Rights.

## Results

### Self-Other Blurring

We compared IOS Scale scores across the Synchronous Movement and Unpredictable groups. IOS scores were not normally distributed according to the Shapiro-Wilk test (W = 0.95, p = .056). Hence, we report Wilcoxon signed-rank test values for this effect. Results showed that IOS scores were significantly higher for the Synchronous Movement group (M = 4.6, SD = 1.8) as compared to the Unpredictable Movement group (M= 2.6, SD = 1.4) (W = 884.5, p < .001) in a directed t-test (Synchronous Movement > Unpredictable Movement) indicating that the IOS scores were significantly higher for the synchronous movement group.

### IAT: D measure

The *D* score (*M* = 0.189, SD = 0.349) was significantly greater than zero (*t_(53)_*_=_ 3.97, *p* < .001, 95% CI [0.253, 0.825]), indicating that the participants tended to react faster to compatible pairings than to incompatible pairings. This result suggested that images of the synchronous movement group were associated with self-words more strongly than with other-words and that images of the unpredictable movement group were associated with other-words more strongly than with self-words.

### IAT: Error Rate

We compared the error rate across the compatible and incompatible blocks. Error rates were not normally distributed according to the Shapiro-Wilk test (W = 0.783 p < .001). Hence, we report Wilcoxon signed rank test values for this effect. Results showed that error rates were lower in the compatible block (M = 4.6, SD = 9) as compared to the incompatible block (M= 5.1, SD = 7.2), but this effect was not significant (W = 478, p =0.131) in a directed t-test (Compatible < Incompatible).

### Control

To establish whether the participants felt more control over their own avatar as compared to the synchronised and unpredictable movement group, we asked participants how much control they felt (see Fig. 5). We conducted a one-way repeated measures ANOVA to assess how much control participants felt over their Avatar, the synchronous group, and unpredictable movement group. Our results indicated a main effect for control (F(2) = 123, p < .001). Subsequent post-hoc tests revealed significant contrasts. Specifically, we found a significant difference between the Avatar Control (M = 3.96, SD = 1.11) and Sync (M = 3.6, SD = 1.1) with higher control reported for the Avatar (Mean Difference = 0.33, SE = 0.165, p_(holm)_ = 0.046). Additionally, there was a significant difference between the Avatar and Unpredict Control (M = 1.57, SD = 0.79), with higher control reported for the Avatar (Mean Difference = 2.38, SE = 0.165, p_(holm)_ < .001). Moreover, a significant difference was observed between the Sync and Unpredict control (Mean Difference = 2.06, SE = 0.165, p_(holm)_ < .001).

### Correlation between Implicit and Explicit Measures

To establish whether there was a relationship between self-other blurring and D measure, we conducted a correlation analysis between IOS difference and D measure across participants. We were unable to find a significant correlation between them when data was collated across conditions (r = -0.069, 95% CI [-0.331, 0.202], p = 0.620).

Additionally, there was no significant relationship between IOS difference and control difference (r = 0.105, 95% CI [−0.167, 0.363], p = 0.448). However, a significant negative correlation was observed between the D measure and control difference (r = −0.318, 95% CI [−0.540, −0.055], p = 0.019), suggesting that greater differences in perceived control were associated with lower D measure scores. This suggests that the greater implicit self-other blurring is associated with lower differences in perceived control between synchrony and unpredictable movement conditions.

## Interim Discussion

The results of Experiment 2 indicate a significant effect of synchronous movement on explicit self–other blurring, as reflected in higher IOS scores for participants in the Synchronous Movement condition compared to those in the Unpredictable Movement condition. The IAT D-score further revealed a strong implicit association between the self and the synchronous movement group, with participants responding more quickly to compatible pairings (self + synchronous group, other + unpredictable movement group). In a subsequent experiment, we introduced a traditional asynchronous control condition (see Fig. 2, third cell) to address concerns arising from the design of Experiment 2. Specifically, the unpredictable movement control combined both temporal jitter and variation in movement types, creating ambiguity about whether the effects were driven by synchrony or by shared behavioral patterns. By employing a standard asynchronous movement control, we were able to benchmark our manipulation against established paradigms in the existing literature, thereby enhancing the interpretability and comparability of our findings within the broader field.

### Experiment 3

#### Methods

Methods and the data analysis plan were pre-registered (https://aspredicted.org/79W_91X). The methods used were identical to Experiment 1 and 2 but the control condition was switched to a group that showed asynchronous movement. We determined our sample size in advance of any data collection, and we report all data exclusions, all manipulations, and all measures in the study. The experimental procedures were approved by the ethical review board at the Institute of Psychology, Humboldt University of Berlin and were conducted in accordance with the American Psychological Association’s Ethical Principles in the Conduct of Research with Human Participants.

#### Participants

We collected the data of 100 participants (30% female, 1% other, M_age_ = 33.880, SD_age_ = 9.37) from Prolific. To acquire exactly 100 participants, we pre-registered our intention to replace participants excluded via the pre-registered subject exclusion criteria.

**Figure 5.**
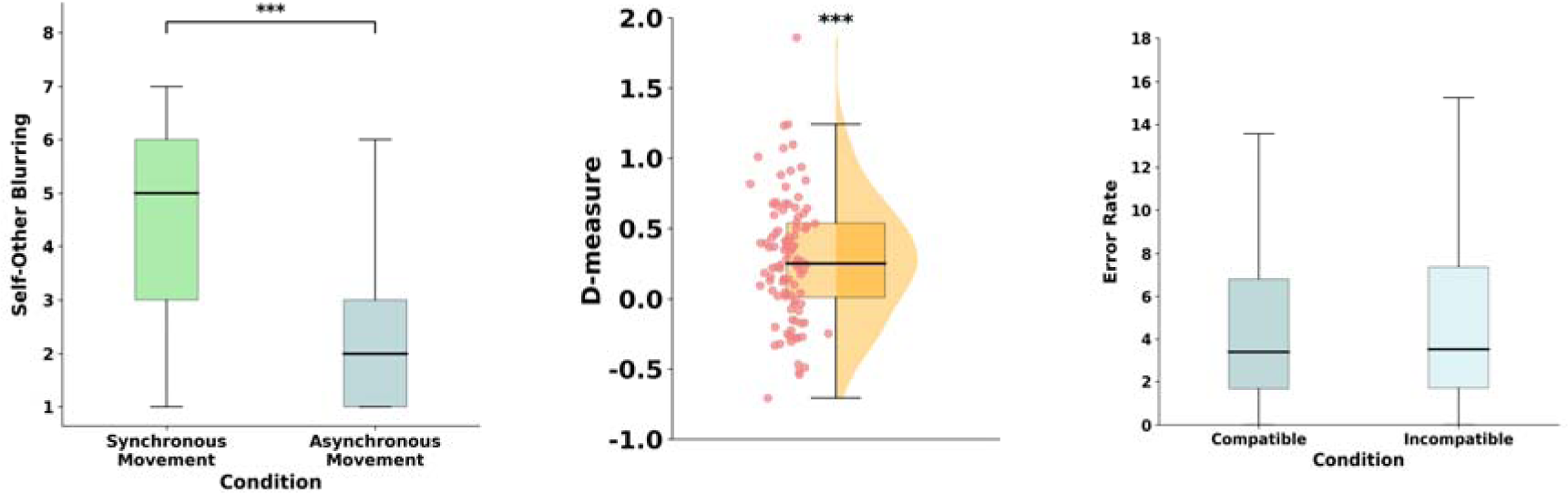
Comparison of IOS Scores for Synchrony vs. Asynchronous Movement Group. Cell 1: Inclusion of Other in the Self (IOS) scores for the Synchrony and Unpredictable Movement groups. Cell 2: Implicit Association Test (IAT) D measure between the Synchrony and Unpredictable Movement groups. Cell 3: Error rates for the Synchrony and Unpredictable Movement groups.

All participants were naïve to the aims of the experiment and compensated at a rate of 6 pounds/hour for their time. Participants provided informed consent before they began the experiment. The experiment was approved by the ethical review board at the Institute of Psychology, Humboldt University of Berlin and was conducted in accordance with the directives laid out in the Helsinki Declaration of Human Rights.

## Results

### Self-Other Blurring

We compared IOS Scale scores across the Synchronous Movement and Asynchronous (Jittered) Movement groups. IOS scores were not normally distributed according to the Shapiro-Wilk test (W = 0.945, p < .001). Hence, we report Wilcoxon signed-rank test values for this effect. Results showed that IOS scores were significantly higher for the Synchronous Movement group (M = 4.68, SD = 1.869) as compared to the Asynchronous Movement group (M = 2.28, SD = 1.615) (W = 3348.000, p < .001) in a directed t-test (Synchronous Movement > Asynchronous Movement), indicating that the IOS scores were significantly higher for the Synchronous Movement group.

### IAT: D measure

The D measure score (M = 0.279, SD = 0.440) was significantly greater than zero (t(99) = 6.345, p < .001, Cohen’s d = 0.635), indicating that participants tended to react faster to compatible pairings than to incompatible pairings. This result suggests that images of the synchronous movement group were associated with self-words more strongly than with other-words, while images of the asynchronous movement group were associated with other-words more strongly than with self-words.

### IAT: Error Rate

We compared the error rate across the compatible and incompatible conditions. Error rates were not normally distributed according to the Shapiro-Wilk test (W = 0.962, p = .006). Hence, we report Wilcoxon signed-rank test values for this effect. Results showed that error rates were lower in the compatible condition (M = 4.307, SD = 4.087) as compared to the incompatible condition (M = 5.156, SD = 4.806), but this effect was not significant (W = 1841.000, z = -1.476, p = 0.07) in a directed t-test (Compatible < Incompatible).

### Sense of Control

To establish whether the participants felt more control over their own avatar compared to the synchronised and asynchronous movement groups, we asked participants how much control they felt. We conducted a one-way repeated measures ANOVA to assess how much control participants felt over their Avatar, the Synchronous group, and Asynchronous group. Our results indicated a main effect for control (F(2) = 131.965, p < .001).

Subsequent post-hoc tests revealed significant contrasts. Specifically, we found a significant difference between the Avatar Control (M = 3.97, SD = 1.21) and Sync (M = 3.33, SD = 1.29) with higher control reported for the Avatar (Mean Difference = 0.640, SE = 0.145, p_(holm)_ < .001). Additionally, there was a significant difference between the Avatar and Asynchronous Control (M = 1.68, SD = 0.875), with higher control reported for the Avatar (Mean Difference = 2.29, SE = 0.151, p_(holm)_ < .001). Moreover, a significant difference was observed between the Sync and Asynchronous control (Mean Difference = 1.65, SE = 0.141, p_(holm)_ < .001).

**Figure 6.**
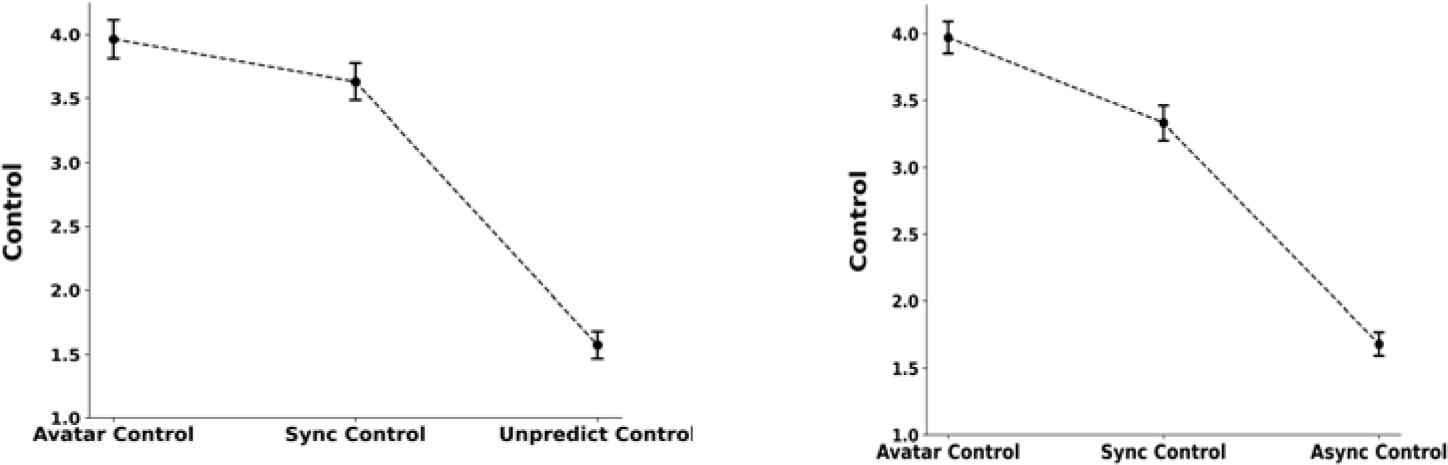
Comparison of Self-Reported Control for Avatar, Synchrony & Unpredictable Movement Group (left cell) and Comparison of Self-Reported Control for Avatar, Synchrony & Asynchronous Movement Group (right cell)

### Correlation between Implicit and Explicit Measures

To establish whether there was a relationship between self-other blurring and D measure, we conducted a correlation analysis between IOS difference and D measure across participants. We found a significant positive correlation between them when data was collated across conditions (r = 0.260, 95% CI [0.067, 0.434], p = 0.009), suggesting that as the IOS difference increases, the D measure also tends to increase.

There was also a marginal correlation between IOS difference and control difference (r = 0.194, 95% CI [−0.002, 0.376], p = 0.053), suggesting a possible but not statistically significant trend toward a relationship between perceived closeness and changes in control. In contrast, the correlation between control difference and the D measure was not significant (r = 0.053, 95% CI [−0.145, 0.247], p = 0.601), providing no evidence of a relationship between control perception and implicit associations in this dataset.

## Discussion

The present series of experiments investigated the impact of interpersonal synchrony on both explicit and implicit measures of self-other blurring. By integrating the Implicit Association Task (IAT) with traditional explicit evaluations using the Inclusion of Other in Self (IOS) scale, we aimed to provide a comprehensive understanding of how synchronous movement influences self-concepts. Across three experiments, participants who engaged in synchronous movement with virtual agents exhibited higher levels of self-other blurring, as evidenced by increased IOS scores and IAT D measures. Higher IAT D measures indicate that they showed faster reaction times and lower error rates in compatible blocks when synchronous movement was paired with self-related words and movement from the control conditions was paired with other-related words, as compared to the incompatible blocks with the reverse pairings. These effects were consistently observed when synchronous movement was contrasted with no movement (Experiment 1), unpredictable movement (Experiment 2), and asynchronous movement (Experiment 3). This means that through their implicit behaviour we can infer that they semantically associated the synchrony group with their self-concept and the control groups with the other concept. This finding bolsters the idea that interpersonal synchrony creates cohesive groups through self-other blurring (Launay et al., 2016). Strengthened implicit associations suggest that participants internalised the synchronous group as part of their self-concept, supporting the notion of self-other blurring at an unconscious level. The significant IAT D measures across all experiments suggest that synchrony strengthens the implicit association between the self and the synchronous group. This implies that synchrony affects not only conscious attitudes but also automatic, unconscious associations. By adapting the IAT to assess self-other associations with specific groups, we introduced a novel application of this implicit measure. The self-other IAT mitigates some limitations of explicit measures, such as social desirability bias and demand characteristics, providing a more robust assessment of unconscious self-concept changes. This methodological innovation can be applied in future research exploring other social phenomena that involve implicit self-concept alterations.

Our use of both the IOS and IAT was motivated by the need to capture distinct but complementary aspects of self–other overlap. The IOS reflects a consciously experienced, embodied sense of affiliation, while the IAT taps into more automatic, semantic-level associations. Together, these tools offer insight into how interpersonal synchrony can affect both felt closeness and the cognitive architecture of the self. The partial convergence of the measures across experiments suggests that synchrony may influence both layers of identity—but not always in lockstep, highlighting the complexity of self-concept change. Furthermore, the inclusion of the IAT helps reduce concerns about demand effects inherent in explicit measures like the IOS.

Our findings on interpersonal synchrony provided new insights into the ongoing debate regarding the distinct yet interconnected roles of implicit and explicit attitudes in social cognition (Fazio & Olson, 2003; Gawronski, 2019; Greenwald & Banaji, 1995). Specifically, it demonstrates that interpersonal synchrony can influence not only explicit experiences of self– other blurring but also implicit self–other representations. This challenges the traditional view that implicit evaluations are slow to change and resistant to brief interventions (Petty et al., 1997), or that they can only be affected by invoking long-standing group memberships (Rydell & McConnell, 2006). In contrast, our findings suggest that implicit self-constructs can shift through even brief, synchrony-based group interactions. This adds to the growing body of research showing that minimally assigned groups in laboratory settings can elicit implicit intergroup bias favouring in-group members (Otten & Moskowitz, 2000; Xiao & Van Bavel, 2019), indicating that implicit evaluations are more flexible than previously assumed and can be shaped by dynamic, short-term social contexts (Gawronski, 2018). Even brief synchronous interactions can rapidly alter implicit self-associations, highlighting the potency of synchrony in shaping social cognition.

Due to our mixed results regarding the relationship between explicit and implicit self-other blurring, we cannot conclusively draw a conclusion from these findings which require more investigation. We found a significant positive correlation between implicit and explicit measures was observed in Experiment 3, suggesting a convergent validity of the two measurement approaches under certain conditions. The lack of a significant correlation between IOS scores and IAT D measures in Experiments 1 and 2, but finding the correlation in Experiment 3, may indicate that the relationship between implicit and explicit self-concept changes depends on the nature of the control condition. When asynchronous movement—a condition more directly contrasting with synchrony—is used, the alignment between implicit and explicit measures becomes more pronounced.

It is not uncommon for IAT/implicit estimates to diverge from their explicit counterparts even when ostensibly targeting the same construct. This divergence is often interpreted as evidence that these measures capture different dimensions of the same construct or target distinct psychological aspects (Slabbinck et al., 2012). This finding holds practical significance for the future study of interpersonal synchrony. If our measure is indeed targeting a different construct, it follows that implicit measures of self-other blurring may exhibit different relationships with the downstream effects typically associated with interpersonal synchrony-induced self-other blurring. Prompting a re-evaluation of the effects commonly linked with explicit self-other blurring, such as willingness to sacrifice for the group (Swann et al., 2010; Whitehouse & Lanman, 2014), threat perception (Swann et al., 2009), and physiological arousal (Swann et al., 2010). Our results support the notion that implicit and explicit evaluations capture different facets of psychological constructs (Gawronski, 2018) and that both are important for understanding the full impact of interpersonal synchrony on self-concept.

In Experiment 2 and 3, participants reported the degree of control they felt over their avatar, the synchronous group, and the unpredictable movement group. In line with Reddish et al.’s (2020) findings, we observed that participants reported higher levels of control for the synchronous group compared to the unpredictable and asynchronous movement group. Feelings of heightened control over the group are known to drive some of the effects associated with synchronous movement (Reddish et al., 2020), thus it is difficult to disentangle feelings of control from effects associated with synchrony (e.g. self-other blurring). With respect to control, even though their avatar and the sync group performed the same action, participants reported the highest level of control for their avatar, indicating that they embodied the avatar.

It is also important to note that correlations between perceived control and self–other blurring measures suggest that control is not straightforwardly linked to either explicit or implicit self–other overlap. In Experiment 2, a significant negative correlation was found between control difference and the IAT D measure, indicating that greater implicit self-blurring with the synchrony group was associated with smaller differences in perceived control ratings between the synchrony and unpredictable groups. This finding aligns with recent suggestions that synchronous movement can lead to a dilution of agency (Jenkins, 2021; Reddish et al., 2020; Zapparoli et al., 2024). As self-other boundaries become blurred during joint action, it becomes harder to identify individual contributions, which may explain the correlation between reduced sense of agency and greater implicit self-other blurring observed here. However, this observation should be interpreted in light of the absence of any other significant correlations with control ratings—neither with explicit self-other blurring in Experiment 2, nor with explicit or implicit self-other blurring in Experiment 3 Together, these findings suggest that while agency may play a role in synchrony effects, its relationship with self–other blurring is not linear or may depend on the contextual factors like the control condition. Further investigation is needed to address how perceived control relates to self-other blurring.

Our study also demonstrates the feasibility and value of using virtual avatars and online platforms to study complex social phenomena like synchrony and self-concept. Although avatar-mediated synchrony lacks the full sensorimotor involvement characteristic of in-person paradigms, it appears to evoke similar psychological outcomes—particularly on the Inclusion of Other in the Self (IOS) scale, which remains the most widely adopted metric in the field. While direct comparisons across modalities remain limited in the literature, this consistency suggests that key features of synchrony, such as temporal alignment and shared intentionality, can effectively be replicated in virtual environments. The avatar-mediated synchrony paradigm allowed for controlled manipulation of movement conditions and minimized confounding variables present in face-to-face interactions. We used avatar-mediated synchrony which has previously been demonstrated to induce explicit self-other blurring (Biswas & Brass, 2024). Another innovative aspect of the study is that the virtual set-up of social agents allows us to study many different control conditions. In the three experiments, we compared synchrony with no movement, unpredictable movement and asynchronous movement.

A post hoc comparison of effect sizes (see Table 1, supplementary materials), it was found that the effect size in Experiment 1 (control condition: no movement) is greater than that in Experiment 2 & 3 (control condition: unpredictable and asynchronous movement. Considering the difference in control conditions between these experiments, the presence and quality of movement may indeed contribute to the magnitude of our effects in Experiment 1. This underscores the importance of employing an appropriate control condition when investigating the effects of synchrony. Determining the optimal control condition to contrast synchronous movement remains a point of contention (Rennung & Göritz, 2016).

Several limitations of the present study also warrant consideration. First, the use of virtual avatars and online experiments, while offering control, may not fully replicate the richness of real-world social interactions. The ecological validity of our findings could be enhanced by conducting similar experiments in face-to-face settings or using more immersive virtual reality technologies. There are additional factors to consider when interpreting and applying our findings. Our focus was on examining the role of interpersonal synchrony within a group setting, which led us to position multiple agents in front of our participants. It’s important to acknowledge that effects may vary in dyadic interactions. Furthermore, our experimental design allowed participants to contrast their experiences with one group against the other, potentially yielding different effects when interacting with each group separately.

In the present study, participants indirectly controlled an avatar engaged in simulated synchronous movement, this mediated online platform is useful for systematically not just for manipulating control conditions but also opens possibilities for exploring broader factors that influence and are influenced by interpersonal synchrony. One such direction involves examining how mediated synchrony might mitigate or amplify pre-existing group identities and biases. The use of online or virtual paradigms is particularly conducive to investigating boundary conditions, as they enable precise and scalable manipulation of experiment and group characteristics. For example, Ye and Gawronski (2016) demonstrated that implicit self-associations can be moderated by factors such as ownership, choice, and object valence—suggesting that similar variables like voluntary participation in synchrony may shape its outcomes. Another promising avenue for future research is to replicate this paradigm in immersive virtual reality or in-person settings to examine whether bodily synchrony amplifies these effects. Together, these directions offer fertile ground for understanding the mechanisms and limits of identity change through coordinated movement.

## Acknowledgements and Funding Information

MBi was supported by the German Academic Exchange Service (DAAD; German: Deutscher Akademischer Austauschdienst). Additionally, this study was funded by DAAD, German Research Foundation (Deutsche Forschungsgemeinschaft, DFG) under Germany’s Excellence Strategy-EXC 2002/1, Science of Intelligence (Project Ref.: 390523135) and the Einstein Foundation Berlin. Further thanks to Dr. Michael Galang (protocol advice). The funders had no role in study design, data collection and analysis, decision to publish, or preparation of the manuscript.

## Declarations

The authors have no competing interests to declare that are relevant to the content of this article.

## Author contributions

Manisha Biswas.: Conceptualization, Writing – Original Draft, Writing – Review & Editing. Marcel Brass: Conceptualization, Writing – Review & Editing, Supervision, Funding acquisition.

## Data Availability Statement

The data that support the findings of this study are openly available at the Open Science Framework [https://osf.io/5jzkq/?view_only=edd8935230f34adba8edd5bf79917599].

## References

Agnew, C. R., & Le, B. (2015). 18 Prosocial Behavior in Close Relationships: An Interdependence Approach. The Oxford Handbook of Prosocial Behavior, 362.

Agnew, C. R., Van Lange, P. A. M., Rusbult, C. E., & Langston, C. A. (1998). Cognitive interdependence: Commitment and the mental representation of close relationships. Journal of Personality and Social Psychology, 74(4), 939–954. 10.1037/0022-3514.74.4.939

Andersen, S. M., & Chen, S. (2002). The relational self: An interpersonal social-cognitive theory. Psychological Review, 109(4), 619–645. 10.1037/0033-295X.109.4.619

Aron, A., Aron, E. N., & Smollan, D. (1992). Inclusion of Other in the Self Scale and the structure of interpersonal closeness. Journal of Personality and Social Psychology, 63, 596–612. 10.1037/0022-3514.63.4.596

Aron, A., Aron, E., Tudor, M., & Nelson, G. (1991). Close Relationships as Including Other in the Self. Journal of Personality and Social Psychology, 60, 241–253. 10.1037/0022-3514.60.2.241

Aron, A., & Fraley, B. (1999). Relationship Closeness as Including Other in the Self: Cognitive Underpinnings and Measures. Social Cognition, 17(2), 140–160. 10.1521/soco.1999.17.2.140

Aron, A., Lewandowski Jr, G. W., Mashek, D., & Aron, E. N. (2013). The self-expansion model of motivation and cognition in close relationships. The Oxford Handbook of Close Relationships, 90–115.

Asendorpf, J. B., Banse, R., & Mücke, D. (2002). Double dissociation between implicit and explicit personality self-concept: The case of shy behavior. Journal of Personality and Social Psychology, 83(2), 380–393. 10.1037/0022-3514.83.2.380

Bargh, J. A., Chaiken, S., Govender, R., & Pratto, F. (1992). The generality of the automatic attitude activation effect. Journal of Personality and Social Psychology, 62(6), 893– 912. 10.1037/0022-3514.62.6.893

Biswas, M., & Brass, M. (2024). Syncing online: A methodological investigation into movement synchrony, proxemics, and self-other blurring in virtual spaces. PLOS ONE, 19(10), e0308843. 10.1371/journal.pone.0308843

Briñol, P., Petty, R. E., & Wheeler, S. C. (2006). Discrepancies between explicit and implicit self-concepts: Consequences for information processing. Journal of Personality and Social Psychology, 91(1), 154–170. 10.1037/0022-3514.91.1.154

Brookes, J., Warburton, M., Alghadier, M., Mon-Williams, M., & Mushtaq, F. (2020). Studying human behavior with virtual reality: The Unity Experiment Framework. Behavior Research Methods, 52(2), 455–463. 10.3758/s13428-019-01242-0

Cirelli, L. K. (2018). How interpersonal synchrony facilitates early prosocial behavior. Current Opinion in Psychology, 20, 35–39. 10.1016/j.copsyc.2017.08.009

Cirelli, L. K., Wan, S. J., & Trainor, L. J. (2016). Social Effects of Movement Synchrony: Increased Infant Helpfulness only Transfers to Affiliates of Synchronously Moving Partners. Infancy, 21(6), 807–821. 10.1111/infa.12140

Collins, R. (2004). Interaction ritual chains. Princeton University Press.

Decety, J., & Sommerville, J. A. (2003). Shared representations between self and other: A social cognitive neuroscience view. Trends in Cognitive Sciences, 7(12), 527–533. 10.1016/j.tics.2003.10.004

Dovidio, J. F., Kawakami, K., & Gaertner, S. L. (2002). Implicit and explicit prejudice and interracial interaction. Journal of Personality and Social Psychology, 82(1), 62–68. 10.1037/0022-3514.82.1.62

Durkheim, E., & Fields, K. E. (1995). The elementary forms of religious life. Free Press.

Fazio, R. H., & Olson, M. A. (2003). Implicit Measures in Social Cognition Research: Their Meaning and Use. Annual Review of Psychology, 54(1), 297–327. 10.1146/annurev.psych.54.101601.145225

Friese, M., Hofmann, W., & Schmitt, M. (2008). When and why do implicit measures predict behaviour? Empirical evidence for the moderating role of opportunity, motivation, and process reliance. European Review of Social Psychology, 19(1), 285–338. 10.1080/10463280802556958

Galdi, S., Arcuri, L., & Gawronski, B. (2008). Automatic Mental Associations Predict Future Choices of Undecided Decision-Makers. Science, 321(5892), 1100–1102. 10.1126/science.1160769

Gawronski, B. (2018). Attitudes and the Implicit-Explicit Dualism. In The Handbook of Attitudes, Volume 1: Basic Principles (2nd ed.). Routledge.

Gawronski, B. (2019). Six Lessons for a Cogent Science of Implicit Bias and Its Criticism. Perspectives on Psychological Science, 14(4), 574–595. 10.1177/1745691619826015

Gómez, Á., Brooks, M. L., Buhrmester, M. D., Vázquez, A., Jetten, J., & Swann, W. B. (2011). On the nature of identity fusion: Insights into the construct and a new measure. Journal of Personality and Social Psychology, 100(5), 918–933. 10.1037/a0022642

Gordon, C. L., & Luo, S. (2011). The Personal Expansion Questionnaire: Measuring one’s tendency to expand through novelty and augmentation. Personality and Individual Differences, 51(2), 89–94. 10.1016/j.paid.2011.03.015

Greenwald, A. G., & Banaji, M. R. (1995). Implicit social cognition: Attitudes, self-esteem, and stereotypes. Psychological Review, 102(1), 4–27. 10.1037/0033-295X.102.1.4

Greenwald, A. G., & Lai, C. K. (2019). Implicit Social Cognition.

Greenwald, A. G., McGhee, D. E., & Schwartz, J. L. (1998). Measuring individual differences in implicit cognition: The implicit association test. Journal of Personality and Social Psychology, 74(6), 1464.

Greenwald, A. G., Nosek, B. A., & Banaji, M. R. (2003). Understanding and using the Implicit Association Test: I. An improved scoring algorithm. Journal of Personality and Social Psychology, 85(2), 197–216. 10.1037/0022-3514.85.2.197

Griffin, D., & Bartholomew, K. (1994). Models of the Self and Other: Fundamental Dimensions Underlying Measures of Adult Attachment. Journal of Personality and Social Psychology, 67, 430–445. 10.1037/0022-3514.67.3.430

Hetts, J. J., Sakuma, M., & Pelham, B. W. (1999). Two Roads to Positive Regard: Implicit and Explicit Self-Evaluation and Culture. Journal of Experimental Social Psychology, 35(6), 512–559. 10.1006/jesp.1999.1391

Hove, M. J., & Risen, J. L. (2009). It’s all in the timing: Interpersonal synchrony increases affiliation. Social Cognition, 27, 949–961. 10.1521/soco.2009.27.6.949

Jenkins, M. (2021). An investigation of “We” agency in co-operative joint actions. Psychological Research.

Jiménez, J., Gómez, Á., Buhrmester, M. D., Vázquez, A., Whitehouse, H., & Swann, W. B. (2016). The Dynamic Identity Fusion Index: A New Continuous Measure of Identity Fusion for Web-Based Questionnaires. Social Science Computer Review, 34(2), 215–228. 10.1177/0894439314566178

Klimmt, C., Hefner, D., Vorderer, P., Roth, C., & Blake, C. (2010). Identification With Video Game Characters as Automatic Shift of Self-Perceptions. Media Psychology, 13(4), 323–338. 10.1080/15213269.2010.524911

Kurdi, B., Seitchik, A. E., Axt, J. R., Carroll, T. J., Karapetyan, A., Kaushik, N., Tomezsko, D., Greenwald, A. G., & Banaji, M. R. (2019). Relationship between the Implicit Association Test and intergroup behavior: A meta-analysis. American Psychologist, 74(5), 569–586. 10.1037/amp0000364

Launay, J., Tarr, B., & Dunbar, R. (2016). Synchrony as an Adaptive Mechanism for LargelJScale Human Social Bonding. Ethology, 122. 10.1111/eth.12528

Markus, H., & Wurf, E. (1987). The Dynamic Self-Concept: A Social Psychological Perspective. Annual Review of Psychology, 38(1), 299–337. 10.1146/annurev.ps.38.020187.001503

Mattingly, B., Mcintyre, K., & Lewandowski Jr, G. (2020). Relationship-Induced Self-concept Change: Theoretical Perspectives and Methodological Approaches (pp. 1– 19). 10.1007/978-3-030-43747-3_1

Mazzurega, M., Pavani, F., Paladino, M. P., & Schubert, T. W. (2011). Self-other bodily merging in the context of synchronous but arbitrary-related multisensory inputs. Experimental Brain Research, 213(2), 213–221. 10.1007/s00221-011-2744-6

McConnell, A. R. (2011). The Multiple Self-Aspects Framework: Self-Concept Representation and Its Implications. Personality and Social Psychology Review, 15(1), 3–27. 10.1177/1088868310371101

Mikulincer, M., Shaver, P. R., & Pereg, D. (2003). Attachment theory and affect regulation: The dynamics, development, and cognitive consequences of attachment-related strategies. Motivation and Emotion, 27(2), 77–102. 10.1023/A:1024515519160

Mogan, R., Fischer, R., & Bulbulia, J. A. (2017). To be in synchrony or not? A meta-analysis of synchrony’s effects on behavior, perception, cognition and affect. Journal of Experimental Social Psychology, 72, 13–20. 10.1016/j.jesp.2017.03.009

Orne, M. T. (1959). The demand characteristics of an experimental design and their implications. *American Psychological Association*, Cincinnati.

Orne, M. T. (2017). On the social psychology of the psychological experiment: With particular reference to demand characteristics and their implications. In Sociological methods (pp. 279–299). Routledge.

Otten, S., & Moskowitz, G. B. (2000). Evidence for Implicit Evaluative In-Group Bias: Affect-Biased Spontaneous Trait Inference in a Minimal Group Paradigm. Journal of Experimental Social Psychology, 36(1), 77–89. 10.1006/jesp.1999.1399

Pacherie, E. (2012). The Phenomenology of Joint Action: Self-Agency vs. Joint-Agency. Joint Attention: New Developments.

Pacherie, E. (2014). How does it feel to act together? Phenomenology and the Cognitive Sciences, 13(1), 25–46. 10.1007/s11097-013-9329-8

Paladino, M.-P., Mazzurega, M., Pavani, F., & Schubert, T. W. (2010). Synchronous Multisensory Stimulation Blurs Self-Other Boundaries. Psychological Science, 21(9), 1202–1207. 10.1177/0956797610379234

Paulhus, D. L. (1984). Two-component models of socially desirable responding. Journal of Personality and Social Psychology, 46(3), 598–609. 10.1037/0022-3514.46.3.598

Perkins, A. W., & Forehand, M. R. (2006). Decomposing the Implicit Self–Concept: The Relative Influence of Semantic Meaning and Valence on Attribute Self–Association. Social Cognition, 24(4), 387–408. 10.1521/soco.2006.24.4.387

Perugini, M., Richetin, J., & Zogmaister, C. (2010). Prediction of behavior. In Handbook of implicit social cognition: Measurement, theory, and applications. (pp. 255–277). The Guilford Press.

Petty, R. E., Wegener, D. T., & Fabrigar, L. R. (1997). Attitudes and attitude change. Annual Review of Psychology, 48(1), 609–647.

Reddish, P., Fischer, R., & Bulbulia, J. (2013). Let’s Dance Together: Synchrony, Shared Intentionality and Cooperation. PLOS ONE, 8(8), e71182. 10.1371/journal.pone.0071182

Reddish, P., Tong, E. M. W., Jong, J., & Whitehouse, H. (2020). Interpersonal synchrony affects performers’ sense of agency. Self and Identity, 19(4), 389–411. 10.1080/15298868.2019.1604427

Rennung, M., & Göritz, A. S. (2016). Prosocial Consequences of Interpersonal Synchrony: A Meta-Analysis. Zeitschrift Für Psychologie, 224(3), 168–189. 10.1027/2151-2604/a000252

Rimé, B., & Páez, D. (2023). Why We Gather: A New Look, Empirically Documented, at Émile Durkheim’s Theory of Collective Assemblies and Collective Effervescence. Perspectives on Psychological Science, 18(6), 1306–1330. 10.1177/17456916221146388

Rusbult, C. E., & Agnew, C. R. (2010). Prosocial motivation and behavior in close relationships. In Prosocial motives, emotions, and behavior: The better angels of our nature (pp. 327–345). American Psychological Association. 10.1037/12061-017

Rydell, R. J., & McConnell, A. R. (2006). Understanding implicit and explicit attitude change: A systems of reasoning analysis. Journal of Personality and Social Psychology, 91(6), 995.

Schimmack, U. (2021). The Implicit Association Test: A Method in Search of a Construct. Perspectives on Psychological Science, 16(2), 396–414. 10.1177/1745691619863798

Schubert, T. W., & Otten, S. (2002). Overlap of Self, Ingroup, and Outgroup: Pictorial Measures of Self-Categorization. Self and Identity, 1(4), 353–376. 10.1080/152988602760328012

Slabbinck, H., De Houwer, J., & Van Kenhove, P. (2012). The Pictorial Attitude Implicit Association Test for need for affiliation. Personality and Individual Differences, 53(7), 838–842. 10.1016/j.paid.2012.06.016

Slotter, E. B., & Gardner, W. L. (2009). Where do you end and I begin? Evidence for anticipatory, motivated self–other integration between relationship partners. Journal of Personality and Social Psychology, 96(6), 1137–1151. 10.1037/a0013882

Slotter, E. B., & Gardner, W. L. (2012a). How Needing You Changes Me: The Influence of Attachment Anxiety on Self-concept Malleability in Romantic Relationships. Self and Identity, 11(3), 386–408. 10.1080/15298868.2011.591538

Slotter, E. B., & Gardner, W. L. (2012b). The dangers of dating the “bad boy” (or girl): When does romantic desire encourage us to take on the negative qualities of potential partners? Journal of Experimental Social Psychology, 48(5), 1173–1178. 10.1016/j.jesp.2012.05.007

Smith, E. R. (1998). Mental representation. The Handbook of Social Psychology, 1, 391.

Smith, E. R. (2008). Embodied Grounding: Social, Cognitive, Affective, and Neuroscientific Approaches (1st ed.). Cambridge University Press. 10.1017/CBO9780511805837

Smith, E. R., Coats, S., & Walling, D. (1999). Overlapping Mental Representations of Self, In-Group, and Partner: Further Response Time Evidence and a Connectionist Model. Personality and Social Psychology Bulletin, 25(7), 873–882. 10.1177/0146167299025007009

Swann Jr., W. B., Gómez, Á., Huici, C., Morales, J. F., & Hixon, J. G. (2010). Identity fusion and self-sacrifice: Arousal as a catalyst of pro-group fighting, dying, and helping behavior. Journal of Personality and Social Psychology, 99(5), 824–841. 10.1037/a0020014

Swann, W. B., Gómez, Á., Dovidio, J. F., Hart, S., & Jetten, J. (2010). Dying and Killing for One’s Group: Identity Fusion Moderates Responses to Intergroup Versions of the Trolley Problem. Psychological Science, 21(8), 1176–1183. 10.1177/0956797610376656

Swann, W. B., Gómez, Á., Seyle, D. C., Morales, J. F., & Huici, C. (2009). Identity fusion: The interplay of personal and social identities in extreme group behavior. Journal of Personality and Social Psychology, 96(5), 995–1011. 10.1037/a0013668

Swann, W. B., Jetten, J., Gómez, Á., Whitehouse, H., & Bastian, B. (2012). When group membership gets personal: A theory of identity fusion. Psychological Review, 119(3), 441–456. 10.1037/a0028589

Swanson, J. E., Swanson, E., & Greenwald, A. G. (2001). Using the Implicit Association Test to investigate attitude-behaviour consistency for stigmatised behaviour. Cognition & Emotion, 15(2), 207–230. 10.1080/02699930125706

Synofzik, M., Vosgerau, G., & Newen, A. (2008). Beyond the comparator model: A multifactorial two-step account of agency. Consciousness and Cognition, 17(1), 219–239. 10.1016/j.concog.2007.03.010

Tajfel, H., Billig, M. G., Bundy, R. P., & Flament, C. (1971). Social categorization and intergroup behaviour. European Journal of Social Psychology, 1(2), 149–178. 10.1002/ejsp.2420010202

Tajfel, H., Turner, J. C., Worchel, S., & Austin, W. G. (1986). Psychology of intergroup relations. Chicago: Nelson-Hall, 7–24.

Tarr, B., Launay, J., Cohen, E., & Dunbar, R. (2015). Synchrony and exertion during dance independently raise pain threshold and encourage social bonding. Biology Letters, 11(10), 20150767. 10.1098/rsbl.2015.0767

Tarr, B., Launay, J., & Dunbar, R. I. M. (2014). Music and social bonding: Âœself-otherâ merging and neurohormonal mechanisms. Frontiers in Psychology, 5. 10.3389/fpsyg.2014.01096

Valdesolo, P., & DeSteno, D. (2011). Synchrony and the social tuning of compassion. Emotion, 11, 262–266. 10.1037/a0021302

Van Lange, P. A. M., Rusbult, C. E., Drigotas, S. M., Arriaga, X. B., Witcher, B. S., & Cox, C. L. (1997). Willingness to sacrifice in close relationships. Journal of Personality and Social Psychology, 72(6), 1373–1395. 10.1037/0022-3514.72.6.1373

Wegner, D. M., Sparrow, B., & Winerman, L. (2004). Vicarious Agency: Experiencing Control Over the Movements of Others. Journal of Personality and Social Psychology, 86(6), 838–848. 10.1037/0022-3514.86.6.838

Whitehouse, H., & Lanman, J. A. (2014). The Ties That Bind Us: Ritual, Fusion, and Identification. Current Anthropology, 55(6), 674–695. 10.1086/678698

Wilson, T. D., Lindsey, S., & Schooler, T. Y. (2000). A model of dual attitudes. Psychological Review, 107(1), 101–126. 10.1037/0033-295X.107.1.101

Xiao, Y. J., & Van Bavel, J. J. (2019). Sudden shifts in social identity swiftly shape implicit evaluation. Journal of Experimental Social Psychology, 83, 55–69. 10.1016/j.jesp.2019.03.005

Zapparoli, L., Mariano, M., Sacheli, L. M., Berni, T., Negrone, C., Toneatto, C., & Paulesu, E. (2024). Self-other distinction modulates the sense of self-agency during joint actions. Scientific Reports, 14(1), 30055. 10.1038/s41598-024-80880-7

