## SupplementaryMaterials for "Interpersonal Synchrony and Implicit Identification"

**Supplementary Materials**

**Effect Size Comparison**

| **Table 1. Comparison of effect sizes and correlation across three experiments** | | | | | |  |
| --- | --- | --- | --- | --- | --- | --- |
|  | **Comparison** | **IOS Effect Size** | **D measure Effect Size** | **Error Rate Effect Size** | **IOS - D Measure Correlation** | **Control Effect Size Difference** |
| **Experiment 1** | *Synchronous vs. No Movement* | 0.970*** | 0.902*** | -0.299* | 0.054 | NA |
| **Experiment 2** | *Synchronous vs. Unpredictable* | 0.787*** | 0.541*** | -0.186 | -0.069 | 2.051*** |
| **Experiment 3** | *Synchronous vs. Asynchronous* | 0.832*** | 0.635*** | -0.175 | 0.260** | 1.045*** |

**Exclusion Criteria**

Subjects Exclusion Criteria:
1. Attention Check Question: Participants providing incorrect responses to the attention check question will be excluded from the analysis. This criterion ensures that participants are attentively engaged in the task and not providing random responses.
2. Missing Catch Trials: Participants who are missing more than one catch trial in the synchrony manipulation will be excluded.
3. High IAT Trial Exclusion: Participants will be removed if they have eliminated more than 70% of the IAT trials. This criterion ensures that participants have provided a sufficient number of valid trials for reliable analysis.
4. Low Latency Threshold: Subjects with more than 10% of trials with a latency less than 300 ms will be excluded. This criterion accounts for response speed and identifies participants potentially exhibiting inattentive or random responding.
5. Experiment Duration: Subjects whose total experiment duration exceeds the third quartile (75%)+ 1.5 times the interquartile range (IQR), calculated from the experiment duration of all participants, will be excluded. This criterion helps identify outliers with unusually long durations that may indicate disengagement or lack of attention.
6. IAT Error Rate: Participants with an error rate in the IAT higher than 30% will be excluded. This criterion ensures the removal of participants with a substantial number of inconsistent or inaccurate responses, preserving data quality.
Trial Exclusion Criteria: For individual trial exclusions, the following criteria will be applied:
1. Extreme Latencies: Trials with latencies greater than 10,000 ms or less than 400 ms will be dropped as outliers. These extreme latencies are likely to reflect technical issues or highly unusual response patterns.
2. Z-Score Outlier Criterion: A z-score measure will be employed to identify outliers in reaction time (RT). Trials with z-scores greater than the predetermined threshold of 3 will be removed, ensuring the removal of RT values that deviate significantly from the overall distribution.
